# A power amplification dyad in seahorses

**DOI:** 10.1101/2020.08.13.249466

**Authors:** Corrine Avidan, Steven W Day, Roi Holzman

## Abstract

Ubiquitous constraints derived from the muscle’s structure limit the power capacity of fast contracting muscles. Correspondingly, organisms evolved elastic elements that store energy which, when released, can be used to rapidly accelerate appendages. Such latch-mediated spring actuation (LaMSA) systems comprise of a single elastic element and are used to actuate a single mass. Here we reveal a dual LaMSA system in seahorses, in which two elastic elements actuate two masses: the head as they rapidly swing it towards the prey, and the water mass sucked into the mouth to prevent the prey from escaping. This power-amplified system enhances the speeds of both head rotation and suction flows by x10 compared to similarly-sized fish. Furthermore, the dual system provides temporal coordination between head rotation and suction flows, a novel function for LaMSA. These findings extend the known function, capacity and design of LaMSA systems.

---

Organismal performance is often limited by muscle power, and this is specifically true for fast, explosive motions. This limitation stems from an innate trade-off between muscle force and contraction speed. Because of the mechanics of muscle sarcomeres, muscles can be either forceful or contract fast, but cannot achieve both functions simultaneously (*1, 2*). Latch-mediated spring actuation (LaMSA) systems, in which slow forceful muscles are used to load an elastic element that is kept latched until it is allowed to abruptly release, have repeatedly evolved in many organisms and overcome muscle power limitations (*3*). LaMSA systems enable some of the fastest animal activities, powering the movements of jumping legs in fleas (*4, 5*), the shell-breaking appendages of mantis shrimps (*6, 7*), and the closing jaws in trap-jaw ants (*8*). LaMSA systems are especially common in arthropods, which utilize their deformable chitinous cuticle to store elastic energy and control its release (*9*). Although, variations of LaMSA systems have been described in dozens of invertebrates, in vertebrates it is extremely rare. The ballistic tongues of anurans and chameleons, as well as the jumping legs in frogs, are among the few LaMSA systems known in vertebrates.

As one of the rare vertebrate examples, a LaMSA system has evolved within the Syngnathiformes (*3, 10*), the order containing the power-amplified seahorses, pipefishes, and snipefishes. Fish within these families capture prey using “pivot feeding”, a behavior comprising an abrupt upward rotation of the head towards the prey. Pivot feeding fishes can open their mouth and elevate their head within a few milliseconds, reaching angular speeds of 200 rad s^−1^. Head rotation is powered by the rapid recoil of the epaxial tendon, which is loaded by the epaxial muscles (*10, 11*). It is thought that the head is maintained in the loaded position by a semi-stable configuration of the hyoid within the four-bar linkage system, and that the retroversion of the hyoid triggers head elevation (Fig 1; Rigid Four-Bar System (*10,12*)). The rapid motion of the elongated snout allows seahorses to specialize in feeding on evasive prey by bringing the mouth close to the prey before it can initiate an escape response. However, bringing the mouth close to the prey is not sufficient for prey capture. To successfully engulf their prey, fish generate a flow of water external to their mouth that carries the prey inside. These “suction flows” persist for a short period and are effective only over small spatial scales (*13*). Therefore, to be effective in capturing evasive prey, suction flows must occur simultaneously with head elevation. The LaMSA system in seahorses, however, was previously thought to affect head rotation exclusively, with unknown consequences for the equally-important suction feeding.

**Fig 1.**
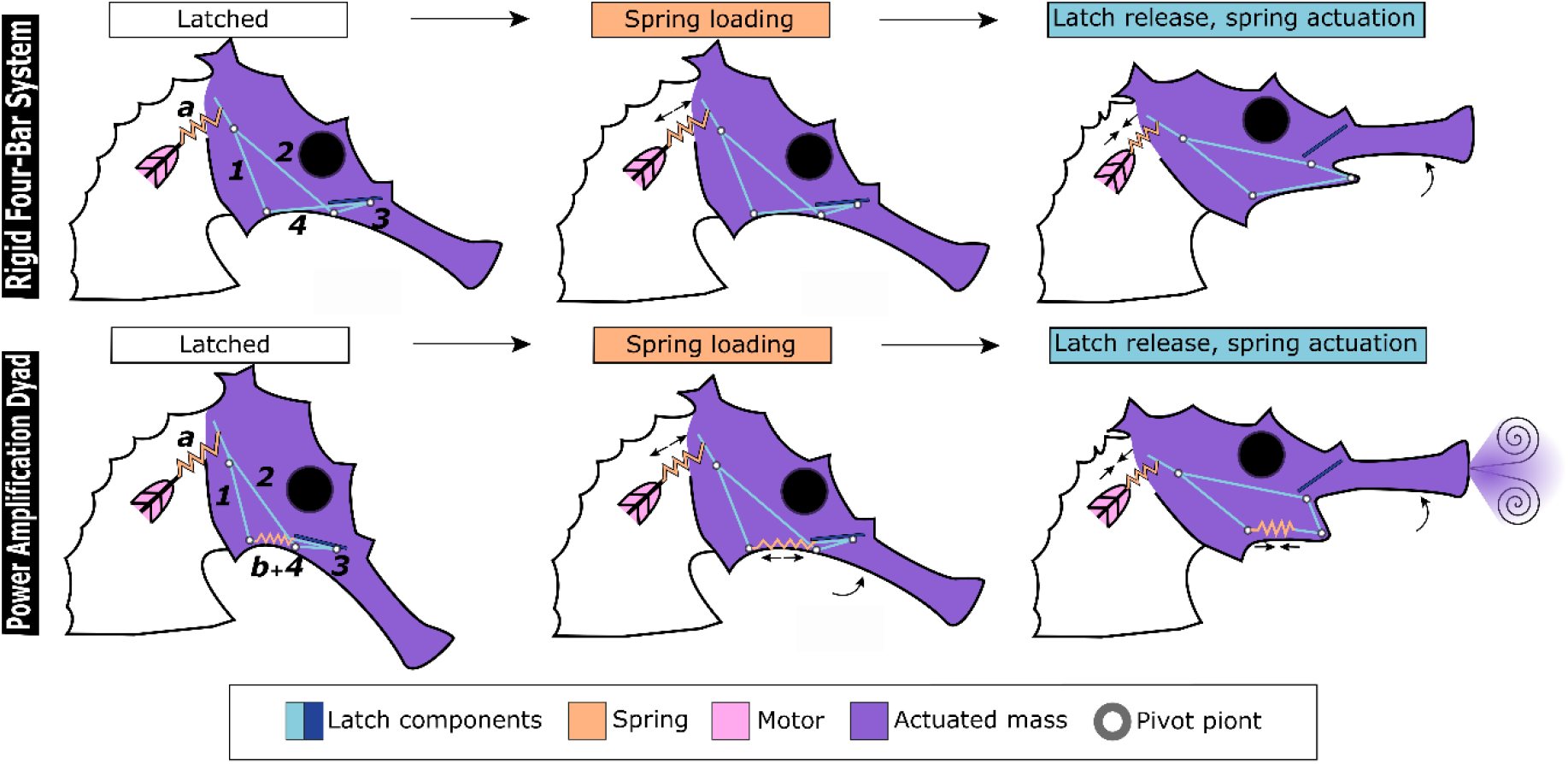
An illustration of the seahorse latch-mediated spring actuation system. The epaxial and sternohyoideus tendons are marked by (a) and (b), respectively, in the leftmost column; 1 is the pectoral girdle, 2 is the neurocranium-suspensorium complex, 3 is the ceratohyal-interhyal complex, and 4 is the urohyal. In a rigid four-bar system configuration (upper row) the urohyal is ossified, resulting in a ridged link, as seen in seahorse relatives (eg. snipefish (*12*), pipefish (*24*)). Accordingly, final hyoid position is set by the observed head elevation and four-bar lengths. In the dyad configuration (bottom row) the presence of the sternohyoideus tendon determines hyoid location. When the seahorse is in the latched and spring loading configuration, the sternohyoideus tendon and the epaxial tendon are loaded and the head cannot be raised. When the latch is released, both tendons recoil to drive rapid head elevation and hyoid retroversion, powering both pivot feeding and suction feeding.

In this study, we characterize suction feeding dynamics, including flow speed, and ask whether seahorses differ from the pattern observed across other fishes. We then employ a novel method to estimate suction power from the external velocity fields and compare this raw power and muscle mass-specific power across fishes. Lastly, we combine material testing and a reconstruction of the hyoid and head movements to calculate the potential for elastic storage of suction power by the sternohyoideus tendon.

We used particle image velocimetry (PIV) to visualize the flow of water external to the mouth during pivot feeding strikes of 12 individuals from three species of seahorses: adult *Hippocampus jayakari, H. fuscus, H. hippocampus*, and *H. jayakari* juvenile (movies <u>S1</u>, <u>S2</u>, <u>S3</u>). Pivot feeding strikes were extremely fast, characterized by time to peak gape of 2.5± 0.18 milliseconds and head rotation speeds of 200 rad s^−1^. These fast kinematics were accompanied by the generation of fast suction flows, which peaked within 2.1 ± 0.011 milliseconds, were coordinated with the time of prey capture, occurred prior to maximal head elevation (Fig 2C) and were an order of magnitude faster than that in other actinopterygians (e.g. bluegill *L. macrochirus*: 33 ± 4 ms (*13*)). Peak measured flow speeds were 0.19 ± 0.004, 0.30 ± 0.016, 0.34 ± 0.007, and 0.32 ± 0.008 m s^−1^ for adult *Hippocampus jayakari, H. fuscus, H. hippocampus* and *H. jayakari juvenile*, respectively. Flow visualization and numerical models revealed that the temporal and spatial dynamics of suction flows are remarkably conserved within actinopterygians. However, the flow speed in seahorses was two orders of magnitude faster than that expected based on the observed relationship between peak gape diameter and flow speed across actinopterygians (Fig 2A; Data from PIV of 14 actinopterygians (*13*)). Similarly to other fish, suction flows in seahorses decayed exponentially with the distance from the center of the mouth, impacting a region of approximately one gape diameter from the mouth (Fig 2B). Taken together, the extreme flow speeds in seahorses and the deviations from the conserved temporal flow dynamics observed across other actinopterygians, indicate that the mechanism generating these flows has been modified to match the snout dynamics, enabling seahorses to suck the prey into the mouth during the pivoting movement of the snout (Fig 2C).

**Fig 2.**
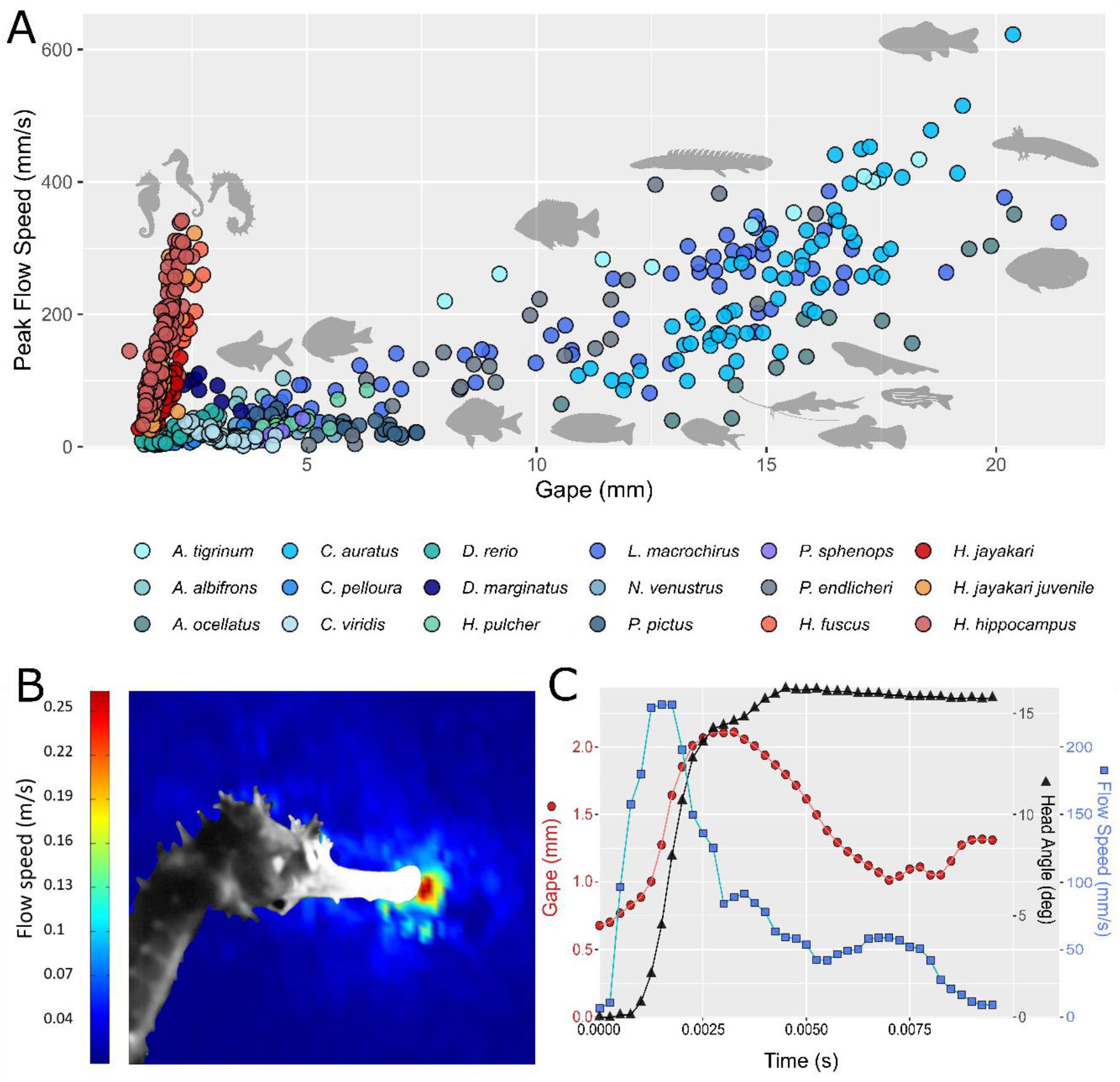
Spatio-temporal patterns of suction feeding flows in seahorses, indicating that a latch-mediated spring actuation system influences these feeding strikes in a couple of ways, without overcoming the hydrodynamic constraints that all actinopterygians face. A) Flow velocities, enhanced by LaMSA, are significantly increased in seahorses compared to other actinopterygians. Colors depict different species, with warm colors depicting species with LaMSA system and cooler colors depicting species without this system. Silhouettes of the represented species are located at approximately peak flow speed positions (Y axis). The presence of a LaMSA system enables seahorses to create much faster flows than expected for their gape size. B) Flow visualization of a suction feeding event, using particle image velocimetry, indicates the conserved spatial patterns of the produced flow, impacting a region of ~ 1 mouth diameter. Image of the studied animal is overlaid a false color image, depicting faster flows as warmer colors and slower flows as colder colors. C) Time series of a typical feeding event, demonstrating that suction flow occurs simultaneously with head rotation and gape opening. The typical time scale of the suction flows in actinopterygians is 0.01-0.02 seconds (*13, 25*).

We hypothesized that the LaMSA system in seahorses is recruited to enhance suction flows and hasten their occurrence. One way of identifying a LaMSA is to compare the power produced by the system with the power capacity of the muscles that drive the system. We calculated the net power used to accelerate the water into the mouth by integrating the suction pressure with respect to buccal volume over time (see SEM Eq S3, S4; (*14*)). We used the Dabiri et al. (*15*) method to estimate the time-varying pressure fields from the PIV flow fields and calculate the pressure at the mouth orifice. We further used the instantaneous flux of water into the mouth to estimate the instantaneous change in buccal volume. These pressure and volume estimates were used to calculate, for each suction feeding PIV video, the total power used to accelerate the water outside of the mouth (hereafter “net suction power”; SEM equation S4). Net suction power was (mean ±SE) 1.63 ± 0.10, 1.56 ± 0.12, 3.76 ± 0.18, and 1.08 ± 0.04 W for adult *Hippocampus jayakari, H. fuscus, H. hippocampus* and *H. jayakari* juvenile, respectively (SEM Fig 1). These power estimates are three orders of magnitude higher than those in other fish with a comparable mouth diameter (0.5-1 mm), and are similar to that of the much larger bluegill *Lepomis macrochirus* and largemouth bass *Micropterus salmoides* (*11*). Average mass-specific net suction power was 3423 ± 223, 3266 ± 310, 4263 ± 207 and 3911 ± 154 W kg^−1^ for adult *Hippocampus jayakari, H. fuscus, H. hippocampus* and *H. jayakari* juvenile, respectively, which is ~3 fold the the maximum recorded from or any vertebrate muscle (1121 W kg^−1^ (*16, 17*)). We repeated the calculation of net suction power using PIV videos of 10 other actinopterygians from reference (*13*), and found that the average mass-specific suction power for these *Hippocampus* species was at least 3 fold higher than those other fishes (3423 ± 350 w kg^−1^ average for seahorses, versus 674 ± 94 w kg^−1^ for *A. ocellatus*, the highest power output measured), and ~10 fold the maximal value measured for bluegill, *Lepomis macrochirus* (321 ± 18 w kg^−1^; this study as well as Camp et al, 2018 (*18*); Fig 3). Species with this LaMSA system use significantly more power during suction feeding (ANOVA, F_(13,146)_ = 95.54, p = <0.001). We therefore conclude that the LaMSA system in seahorses is used to power both pivot feeding and suction feeding, resulting in coordinated, extremely fast head rotation and buccal expansion.

**Fig 3.**
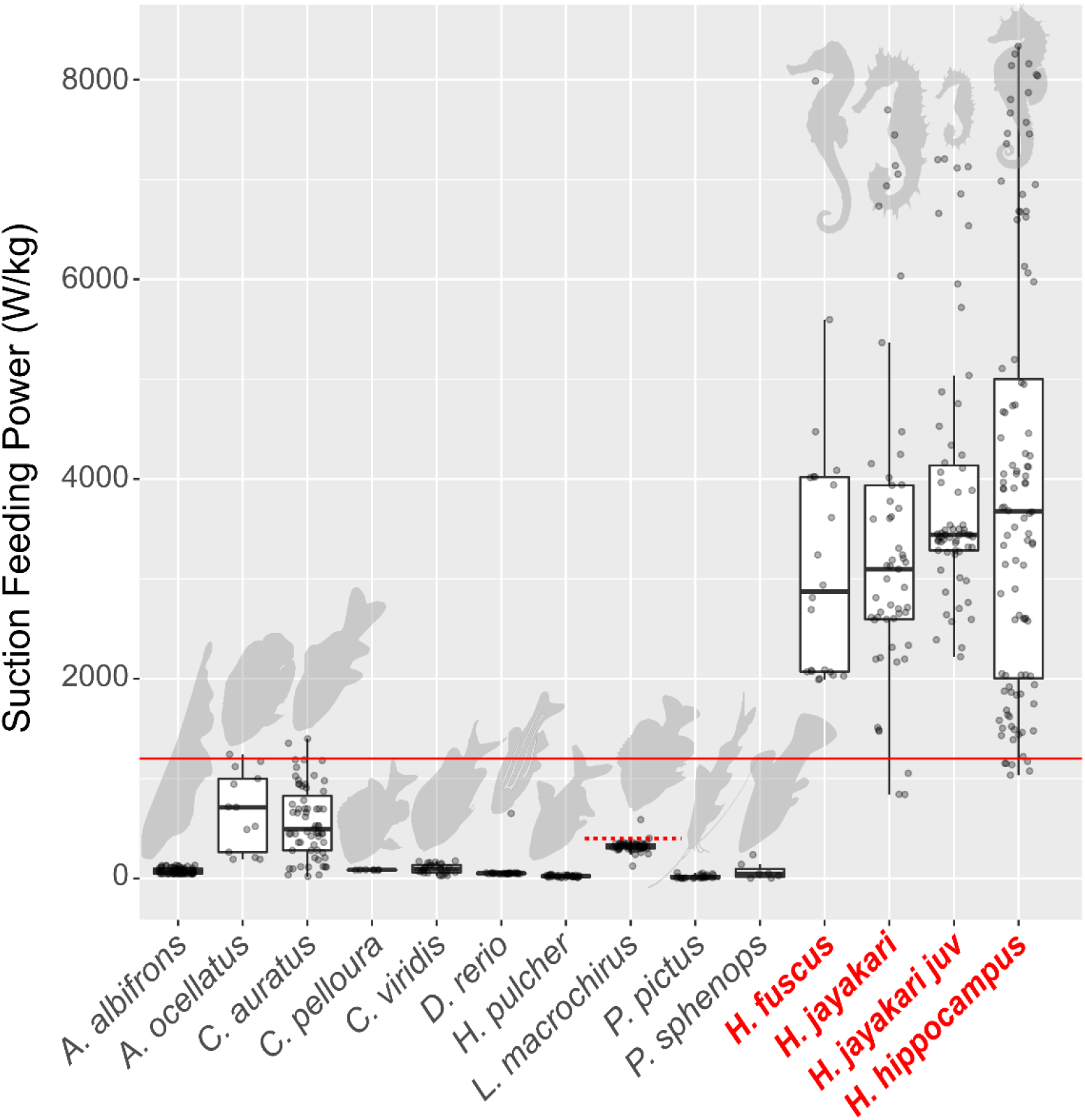
Mass specific power of suction feeding for seahorses (*Hippocampus Fuscus, H. jakari and H. hippocampus*) is X3 times higher the maximum recorded from or any vertebrate muscle (1121 W kg^−1^; solid red line; (*16,17*)) indicating that seahorses use a LaMSA system to power their suction flows. Power for both LaMSA and non-LaMSA fishes (gray circles) was estimated by using the PIV-measured flow field to jointly estimate buccal pressure and suction volume (SEM equation S3), and hence refers to net suction power, i.e. the power used to accelerate the water outside the mouth. Boxes encompass the 2-3^rd^ quartile range, and the horizontal black line is the median estimated power for each species. The dashed line indicates the estimated peak suction feeding power for *L. macrochirus* from Ref (*18*) which agrees with the results from the method used in this study. We therefore conclude that the LaMSA system in seahorses has a dual function, powering both pivot feeding and suction feeding.

In seahorses, head rotation is achieved through the recoil of the epaxial tendon, connecting to the supraoccipital bone at the back of the head (Fig 1; (*19*)). Previous studies have modeled the hyoid movement in pivot feeding Syngnathiformes as part of a four-bar lever system that transmits force and movement from the epaxial tendon (Fig 1; Fig 4A; (*10, 12*)). Buccal expansion is driven by hyoid retroversion, which depresses the floor of the mouth cavity and expands the head laterally. According to this model, suction power comes from the epaxial tendon, i.e. there is a single elastic element which drives these two coordinated functions. However analysis of cleared and stained seahorse specimens revealed that the hyoid is connected to the pectoral girdle by a second tendon (the sternohyoideus tendon); while high-speed kinematic analysis of the four-bar system identified deviations from the movement expected based on a rigid system with the observed geometry. We therefore assert that the transmission of motion and power within a seahorse’s head is more complex than previously described. Specifically, we suggest that the sternohyoideus tendon stores elastic energy that contributes to the generation of suction flows. Indeed, tracking the distal end of the hyoid and the pectoral girdle (the two attachment points of the tendon) revealed that the tendon compresses by up to 55% during hyoid retroversion (Figures 4A and S3). We dissected and measured the force-length relationships of the tendon in 5 individuals and calculated the power input of the compressing tendon to the system, based on the tendon recoil observed from PIV movies (Fig. 4B). A multiple regression mixed-effect model revealed strong correlations between the power contributed by the compressing sternohyoideus tendon and the net suction power (marginal and conditional R^2^= 0.71 and 0.75 respectively, p < 0.001). The recoiling tendon was able to generate more power than used for the estimated net suction power, indicating that the sternohyoideus tendon can power suction feeding by directly pulling on the hyoid. Note that the agreement between two power measurements does not necessarily imply that the epaxial tendon only contributes power to head elevation, because the four-bar lever system (although containing an elastic bar) can still transmit force to the hyoid. Unfortunately, it is not possible to track the dynamics of the epaxial tendon within the PIV setup.

**Fig 4.**
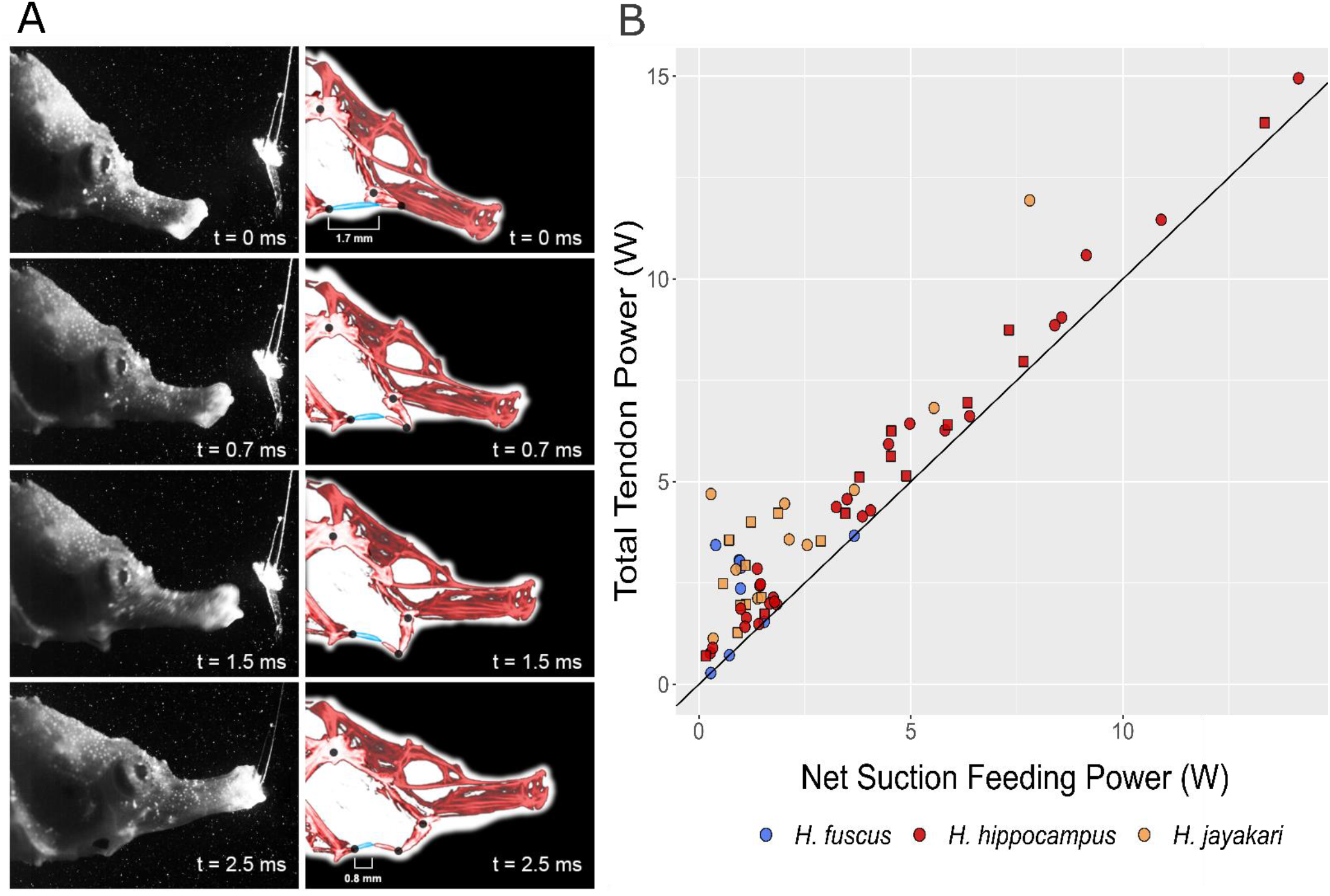
Recoil and power export of the sternohyoideus tendon throughout pivot feeding in seahorses, suggesting that energy is stored within the sternohyoideus tendon, in addition to the previously-recognized epaxial tendon. A) Time series with PIV frames on the left and a CT overlay of the bones (red) and sternohyoideus tendon (blue) on the right. The sternohyoideus tendon shortens by 53% during this strike. All joints of the four bar are represented by black points. B) The power contributed by the recoiling sternohyoideus tendon (SEM eq S5) is significantly correlated with net suction power (marginal and conditional R^2^= 0.71 and 0.75 respectively, p < 0.001), and slightly above the 1:1 line (black), indicating power loss during the strike. The excess power can potentially account for the drag created when moving water through the snout. Power contributed by the compressing sternohyoideus tendon was estimated using the observed contraction of the tendon (from the PIV movies) and the length-force curve of the tendon. Colors represent different species and shapes represent the individuals within each species.

We conclude that the LaMSA system in seahorses represents a unique mechanism in which two functions that are performed simultaneously (i.e. suction feeding and head elevation) are powered by two elastic elements (the epaxial tendon and the sternohyoideus tendon). These elastic elements and the ensuing movements are coordinated, probably by virtue of sharing the same latch system. To the best of our knowledge, the only other known dual-powered LaMSA system is found in mantis shrimps, in which the deformation of the saddle and the rotation of the meral-V store elastic energy that is released for the singular function of accelerating the bashing appendage (*20*). The coordination of head elevation and suction flow is necessary for prey capture because it allows the fish to draw their prey into the mouth before an escape response can be initiated. This is especially important when feeding on Copepods, which can accelerate to escape their predators at up to 300 m s^−2^ (*21*). Myelinated axons help these potential prey to initiate the response within approximately 4 ms from perceiving the hydrodynamic disturbance generated by the moving predatory fish (or its snout; (*22*)). Consequently, delaying the suction flow until after the predator’s mouth has arrived near the prey would have provided the copepods with ample time to escape. With head elevation and suction flow coordination, however, attacks will be successful even though the power-amplification does not break constraints on spatial patterns: i.e. suction flows only affect a region of ~1 gape diameter away from the mouth. In both pipefishes (nested with seahorses within the family Syngnathidae) and species from the sister families Centriscidae (snipefishes) and Fistulariidae (cornetfishes), the hyoid is connected to the pectoral girdle by the ossified urohyal, forming a rigid four-bar lever system (*23*). Although the power amplification dyad in seahorses is likely the derived state within Syngnathidae, to the best of our knowledge, a systematic investigation of urohyal ossification within Syngnathiformes has not been conducted. Pipefishes and snipefishes both use a LaMSA system to power pivot feeding (*10, 12*), and the rapid movement of their hyoid probably implies that suction flows are coordinated with head elevation, similarly to seahorses. This also implies that suction feeding in these ossified species is also power-amplified. However, it is unknown whether suction flows in these species are faster compared with non-amplified flows.

## Animal care

Maintenance and experimental procedures followed the IACUC approved guidelines at the Hebrew University of Jerusalem, overseeing the experiments at IUI Eilat.

## Author Contributions

C.A. and R.H. initiated the project; C.A. conducted the PIV experiments, analyzed and visualized the results. SWD contributed data on the coupled PIV-pressure experiments. All authors contributed to conceptualization and development of the ideas presented, analyses, and interpretation of results. C.A. and R.H. wrote the paper with contribution of SWD. All authors approved the final manuscript.

## Acknowledgments

The study was funded by Israel Science Foundation grant 965/15 to RH. CJ thanks CLiF for financial support. We thank S. Ayalon and B. Eisenberg for providing seahorses for the study. We thank P. Wainwright and S. Longo for comments on the manuscript.

## References

1. S. S. Jahromi, H. L. Atwood, Correlation of structure, speed of contraction, and total tension in fast and slow abdominal muscle fibers of the lobster (*Homarus americanus*). J. Exp. Zool. 171, 25–37 (1969).

2. G. M. Taylor, Maximum force production: Why are crabs so strong? Proc. R. Soc. B Biol. Sci. 267, 1475–1480 (2000).

3. S. J. Longo, S. M. Cox, E. Azizi, M. Ilton, J. P. Olberding, R. St Pierre, S. N. Patek, Beyond power amplification: latch-mediated spring actuation is an emerging framework for the study of diverse elastic systems. J. Exp. Biol. 222, jeb197889 (2019).

4. M. Burrows, How fleas jump. J. Exp. Biol. 212, 2881–2883 (2009).

5. H. C. Bennet-Clark, E. C. Lucey, The jump of the flea: a study of the energetics and a model of the mechanism. J. Exp. Biol. 47, 59–67 (1967).

6. S. N. Patek, W. L. Korff, R. L. Caldwell, Biomechanics: Deadly strike mechanism of a mantis shrimp. Nature. 428, 819–820 (2004).

7. S. N. Patek, B. N. Nowroozi, J. E. Baio, R. L. Caldwell, A. P. Summers, Linkage mechanics and power amplification of the mantis shrimp’s strike. J. Exp. Biol. 210, 3677–3688 (2007).

8. F. J. Larabee, W. Gronenberg, A. V. Suarez, Performance, morphology and control of power-amplified mandibles in the trap-jaw ant Myrmoteras *(Hymenoptera: Formicidae*). J. Exp. Biol. 220, 3062–3071 (2017).

9. J. F. V. Vincent, Arthropod cuticle: A natural composite shell system. Compos. Part A Appl. Sci. Manuf. 33, 1311–1315 (2002).

10. S. Van Wassenbergh, J. A. Strother, B. E. Flammang, L. A. Ferry-Graham, P. Aerts, Extremely fast prey capture in pipefish is powered by elastic recoil. J. R. Soc. Interface. 5, 285–296 (2008).

11. S. Van Wassenbergh, S. W. Day, L. P. Hernández, T. E. Higham, T. Skorczewski, Suction power output and the inertial cost of rotating the neurocranium to generate suction in fish. J. Theor. Biol. 372, 159–167 (2015).

12. S. J. Longo, T. Goodearly, P. C. Wainwright, Extremely fast feeding strikes are powered by elastic recoil in a seahorse relative, the snipefish, *Macroramphosus scolopax*. Proc. R. Soc. B Biol. Sci. 285, 1–8 (2018).

13. C. N. Jacobs, R. Holzman, Conserved spatio-temporal patterns of suction-feeding flows across aquatic vertebrates: a comparative flow visualization study. J. Exp. Biol. 221, 1–11 (2018).

14. A. M. Carroll, P. C. Wainwright, Energetic limitations on suction feeding performance in centrarchid fishes. J. Exp. Biol. 212, 3241–3251 (2009).

15. J. O. Dabiri, S. Bose, B. J. Gemmell, S. P. Colin, J. H. Costello, An algorithm to estimate unsteady and quasi-steady pressure fields from velocity field measurements. J. Exp. Biol. 217, 331–336 (2014).

16. R. M. Alexander, H. C. Bennet-Clark, Storage of elastic strain energy in muscle and other tissues. Nature. 265, 114–117 (1977).

17. R. M. N. Alexander, Tendon elasticity and muscle function. Comp. Biochem. Physiol. – A Mol. Integr. Physiol. 133, 1001–1011 (2002).

18. A. L. Camp, T. J. Roberts, E. L. Brainerd, Bluegill sunfish use high power outputs from axial muscles to generate powerful suction-feeding strikes. J. Exp. Biol. 221, jeb178160 (2018).

19. H. Leysen, J. Christiaens, B. de Kegel, M. N. Boone, L. van Hoorebeke, D. Adriaens, Musculoskeletal structure of the feeding system and implications of snout elongation in *Hippocampus reidi* and *Dunckerocampus dactyliophorus*. J. Fish Biol. 78, 1799–1823 (2011).

20. S. N. Patek, The power of mantis shrimp strikes: Interdisciplinary impacts of an extreme cascade of energy release. Integr. Comp. Biol. 59, 1573–1585 (2019).

21. R. J. Waggett, E. J. Buskey, Calanoid copepod escape behavior in response to a visual predator. Mar. Biol. 150, 599–607 (2007).

22. A. D. Davis, T. M. Weatherby, D. K. Hartline, P. H. Lenz, Myelin-like sheaths in copepod axons. Nature. 398, 571 (1999).

23. G. Roos, H. Leysen, S. Van Wassenbergh, A. Herrel, P. Jacobs, M. Dierick, P. Aerts, D. Adriaens, Linking Morphology and Motion: A Test of a Four-Bar Mechanism in Seahorses. Physiol. Biochem. Zool. 82, 7–19 (2009).

24. B. A. Bergert, P. C. Wainwright, Morphology and kinematics of prey capture in the syngnathid fishes *Hippocampus erectus* and *Syngnathus floridae*. Mar. Biol. 127, 563–570 (1997).

25. S. W. Day, T. E. Higham, R. A. Holzman, S. Van Wassenbergh, Morphology, kinematics, and dynamics: the mechanics of suction feeding in fishes. Integr. Comp. Biol. 55, 21–35 (2015).

