## Supplementary material for "A power amplification dyad in seahorses": Methods

**Supplementary Methods**

We studied three species of seahorses (*Hippocampus jayakari, H. fuscus, H. hippocampus*). For each species, two to five adult individuals were used as well as five *Hippocampus jayakari* juvenile. Flow data were collected using the same methods as in Higham et al. (2005) (reference (*26*)) and Jacobs and Holzman (2018) (reference (*13*)) to facilitate comparisons among species. Seahorse species were collected locally from the Gulf of Aqaba and the Mediterranean Sea. The study animals were housed in indoor aquaria and fed daily with live mysid shrimp. Animal maintenance and experimental procedures followed the IACUC approved guidelines at the Hebrew University in Jerusalem, which oversees the experiments at the Inter-University Institute in Eilat.

*Particle image velocimetry (PIV)*

The principles behind PIV can be found in detail in Stamhuis, E (2006) (reference (*27*)) and Taylor et al. (2014) (reference (*28*)) and are explained here in brief. A solid state continuous wave laser (Coherent Magnum II Laser, 680 nm, 1.2 Watts, 10 degree fan angle, Santa Clara, CA, USA) equipped with a built-in optical system was used to illuminate neutrally buoyant 10 µm hollow glass spheres within the tank. The light sheet created by the laser has a <0.5 mm thickness and ∼5 cm height. Videos of seahorse feeding events were recorded at 4,000 frames $s^{-1}$ and a resolution of 640x640 pixels using a Photron SA3 2000 high-speed video camera (Photron, Tokyo, Japan) equipped with 105 mm Nikon lens ( f=2.8, Nikon, Tokyo, Japan). In all experiments, the camera was positioned orthogonal to the light sheet to capture lateral views of the feeding animal. Distances in the videos were scaled by recording an image of a ruler. Prey were suspended on a thin wire. The study animals were trained to approach the prey from a perch position within the laser sheet to ensure that the light sheet was aligned with the sagittal plane of the head. A commercial video camera (Go-pro Hero4, GoPro Inc., San Mateo, CA, USA) was located above the aquarium to verify the position of the animal's mouth with respect to the light sheet, and only sequences in which the two were aligned were used. Over 250 feeding strikes were analyzed for our three study species (a mean of 25 strikes per individual). High-speed PIV videos of feeding sequences were analyzed using MatPIV, a freely available toolbox for analyzing PIV (*29–31*) in MATLAB (The MathWorks, Natick, MA, USA). MatPIV treats the high-speed video sequences as a series of image pairs, each consisting of two successive frames (at 0.25 ms intervals). For each image pair, MatPIV estimates the flow speed and direction at each location on a regularly spaced grid of 64x64 cells (16×16 pixels each, with 50% overlap between adjacent cells). The algorithm also calculates a signal-to-noise ratio used to validate the velocity measurements.

*Kinematics*

To characterize skull kinematics, we digitized in each frame the location of the proximal tip of the upper and lower jaw, the center of the eye socket, the prey's center of mass, and the joint locations of the four-bar system; the proximal tip of the hyoid bone (when visible), the ventral attachment of the pectoral girdle, hyoid-suspensorium attachment, and the neurocranial pectoral girdle attachment, using DLTdv5 ((*32*); SEM Fig 2). Due to the intense light from the laser sheet, most of these locations were visible as the skin becomes bright and semi translucent. However, a few of the locations from the four-bar system could only be estimated based on the axis of rotation within the videos. From the overall digitization, we calculated for each frame the following variables (*33*): (1) gape diameter, defined as the distance between the upper and lower jaw points; (2)

sternohyoideus tendon length, defined as the distance between the proximal tip of the hyoid bone and the ventral attachment of the pectoral girdle; and (3) head rotation, defined as the rotational angle of the center of the mouth to the neurocranial pectoral girdle attachment. From the time-dependent patterns of gape diameter and head rotation we determined peak excursions and their respective timings. We defined time to peak gape (TTPG) as the time from when gape diameter first exceeded 20% of its maximal value to the time when it first exceeded 95% of its maximal value. Time to peak hyoid displacement was similarly calculated (*33*). While we acknowledge that use of external markers is not as ideal as x-ray moving morphology (XROMM), size limitations prevented that option and the seahorse's uniquely thin skin, paired with the high intensity laser, offer a sufficiently clear resolution of the movement of the bones to track the above points accurately.

*Spatio-temporal patterns*

To examine the spatial distribution of flow velocities in front of the animal's mouth, we extracted flow speeds at a distance of ½ gape diameter from the mouth on the center line. Because, PIV measurements near solid boundaries are prone to bias due to movements of the animal's body interfering with the signal from the moving particles, a greater distance was chosen. The flux of water flowing into the mouth was calculated as the integral, from the time of mouth opening to closing, of flow speed at the mouth aperture multiplied by gape area. For these calculations, flow speed at the mouth aperture was calculated in the animal-bound frame of reference, i.e. as the sum of ram speed and flow speed at the earthbound frame of reference (*34*). This procedure assumes that the magnitude of flow speed is identical across the mouth orifice, an assumption supported by a CFD model of suction flows (*35*) as well as by theory (see discussion in (*36*)). We further assume that the mouth is circular (i.e. non-elliptical).

*Flow power*

We used a method for estimating the pressure field (developed by (*15*)) corresponding to velocity field measurements obtained by using particle image velocimetry. The corresponding pressure field is determined based on median polling of several integration paths through the pressure gradient field, in order to reduce the effect of measurement errors that accumulate along individual integration paths. Integration paths are restricted to the nodes of the measured velocity field, thereby eliminating the need for measurement interpolation during this step. This method is based on direct integration of the pressure gradient term in the Navier–Stokes equation for incompressible flow:

$$\nabla p = -\rho\left(\frac{Du}{Dt} - v\nabla^2 u\right) \qquad (S1)$$

where v is the kinematic viscosity of the fluid. The pressure difference described in Eqn 1 is between the ambient water pressure at far distances from the suction feeding event and the flow within 1 gape diameter of the mouth of the fish. The instantaneous fluid particle acceleration Du/Dt required for calculation of the pressure gradient in Eqn 1 is calculated from a single time-dependent velocity field as:

$$\frac{Du}{Dt} = \frac{\partial u}{\partial t} + (u \cdot \nabla)u \qquad (S2)$$

Eight families of integration paths were used, originating at the domain boundary and propagating toward each grid point. Velocity fields were processed according to Dabiri et al. (2014) (reference (*15*)) with the exceptions that (1) no temporal smoothing was used due to the nature of the suction feeding flows; and (2) the flow values within the snout were artificially replaced with the flow velocities at the mouth orifice. This was done because the above method requires all integration paths to start at a domain boundary, and masking the fish snout creates an unusable boundary. Therefore, the flow speed at the mouth aperture is duplicated, assuming constant velocity within the mouth.

The method was validated using a dataset of simultaneously captured PIV footage and measured intraoral pressure reported in Higham et al. (*37*). In that study, a Millar SPR-407 microcatheter-tipped pressure transducer was positioned flush with the buccal cavity. The sensing element was thereby physically shielded by a plastic cannula, but exposed to pressure changes by a short fluid path in the buccal cavity during the feeding strikes. These strikes were simultaneously assessed using PIV to measure the fluid velocity field during suction feeding. By using the PIV information to calculate the pressure, as explained above, and then comparing it with the measured pressure, we discovered a tight correlation between PIV-estimated and measured buccal pressure (SEM Fig 3). This novel method enabled significant insights into feeding fishes while eliminating the influences of risky surgery or changes in behavior due to attachments on the fish.

Mechanical work (W; Joules) was calculated by numerical integration of pressure *p* at the orifice with respect to the change in volume from the onset of gape opening until the end of suction flow.

$$W = \int_{V_i}^{V_f} p \, dV \qquad (S3)$$

It should be noted that the change in buccal volume (*dV*) was estimated by the instantaneous flux of water into the mouth, calculated from the flow velocities at the mouth orifice multiplied by the assumed radially symmetric opening(*14*). By estimating both pressure and flux throughout the feeding strike we can obtain time-dependent power rather than peak work and power which would rely on maximal buccal volume measured post-mortem(*14*). Work (W) was divided by duration to calculate raw power, P, (Watts) and both values were divided by epaxial muscle mass to estimate mass specific power (Watts $kg^{-1}$).

$$P = \frac{\mathrm{W}}{\Delta t_{AR}} \qquad (S4)$$

Where $t_{AR}$ is the duration from the onset of gape opening until the end of suction flow. Muscle mass measurements incorporate both epaxial and hypaxial muscles, and were taken from specimens in this study when possible. However, for *A. ocellatus* (*38*), *C. auratus* (*39*), and *L. macrochirus* (*14*, *18*), average muscle mass was taken from the literature in order to approximate power.

*Tendon power*

In order to determine the amount of power within the sternohyoideus tendon, we dissected and measured the force-length relationships of the tendon for five individuals. The pectoral girdle and the hyoid were attached, allowing the sternohyoideus, which connects them, to stretch freely. Tendons were kept wet and stretched at 0.01 mm increments and held at each position for 30 s using a custom made material testing apparatus. Each tendon was stretched 0.3 mm past the maximum length measured in the PIV videos. The slope of ligament force as a function of length was used to calculate the force capacity of the ligament (F). The total power contributed by the ligament was estimated as the integral of the force capacity of the ligament (F) multiplied by the change in ligament length ($L_t$) estimated in each time step (SEM fig 4), with respect to time:

$$P = \int_{t_i}^{t_f} F\, L_t \qquad (S5)$$

Where $t_i$ and $t_f$ are the times of initial and final hyoid movements (seconds).

*Statistical analyses*

A mixed-effects model was used to examine the correlation between total tendon power and net suction feeding power, with total tendon power as the fixed factor, with individuals treated as random factors. The mixed-effect model was run using the lme4 package in R (https://www.r-project.org/), and the P-value was obtained by using an ANOVA to compare this model and an intercept-only model. $R^2$ values were calculated using the package MuMin. The marginal $R^2$ value describes the proportion of variance explained solely by the fixed factors and the conditional $R^2$ describes the proportion of variance explained by both the fixed and random factors.

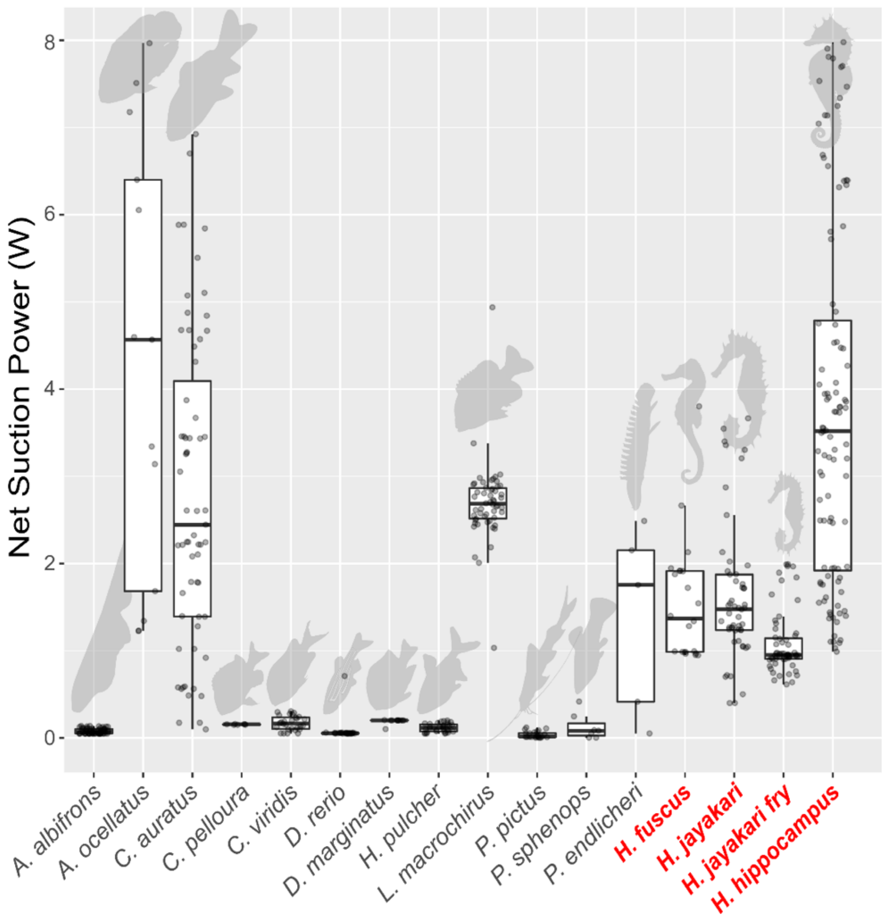


SEM Fig 1. Total net suction power for seahorses is three orders of magnitude higher than that for other fish with a comparable mouth diameter (0.5-1 mm). It is also similar to that of the much larger bluegill *Lepomis macrochirus*, *Carassius auratus*, and *Astronotus ocellatus*. Power for both LaMSA and non-LaMSA fishes (gray circles) was estimated using PIV-measured flow field to jointly estimate buccal pressure and suction volume, and hence refers to net suction power i.e. the power used to accelerate the water outside the mouth. Boxes encompass the 2-3rd quantile range; horizontal black line is the median estimated power for each species.

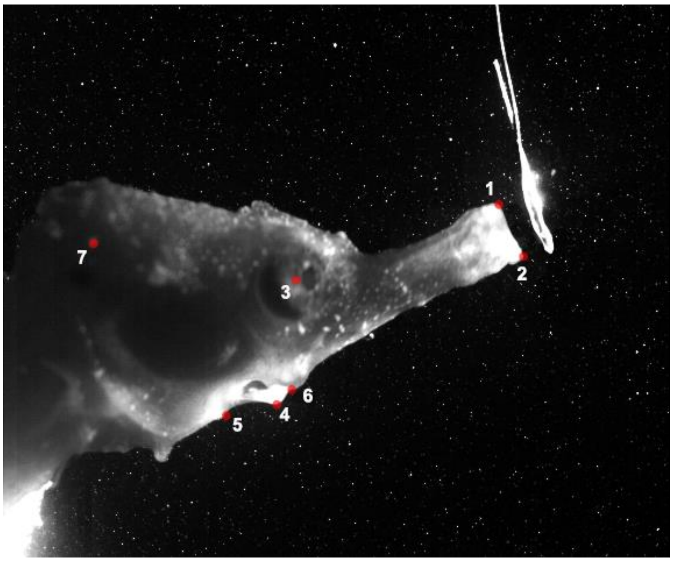


SEM Fig 2. The locations marked for analysis of (1) the proximal tip of the upper and (2) lower jaw, (3) the center of the eye socket, the (not shown) prey's center of mass, and the joint locations of the four-bar system: (4) the proximal tip of the hyoid bone (when visible), (5) the ventral attachment of the pectoral girdle, (6) hyoid-suspensorium attachment, and (6) the neurocranial pectoral girdle attachment. The intense light from the laser sheet causes the skin to become bright and semi-translucent enabling visualization.

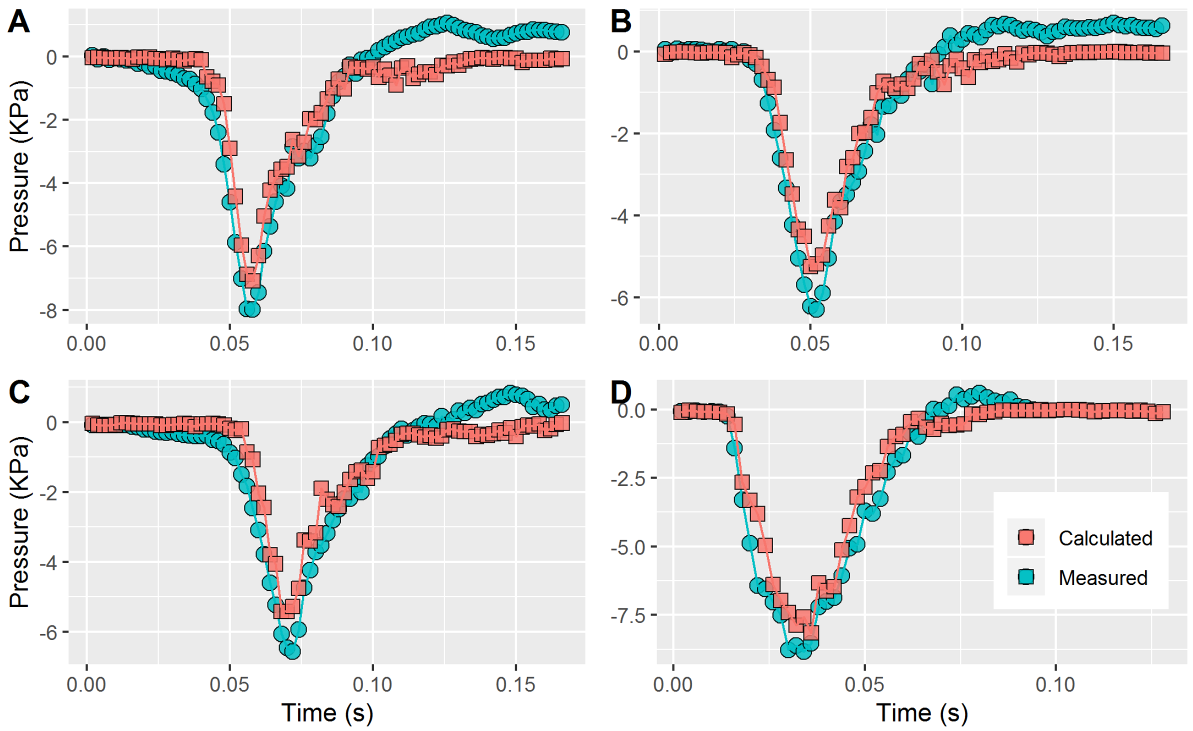


SEM Fig 3. Highly correlated pressures, verifying the method of calculating pressure from PIV footage. Measured pressure data were obtained from a microcatheter-tipped pressure transducer within the buccal cavity of the largemouth bass (*Micropterus salmoides*)(*37*), and simultaneously captured PIV images were used to calculate pressure.

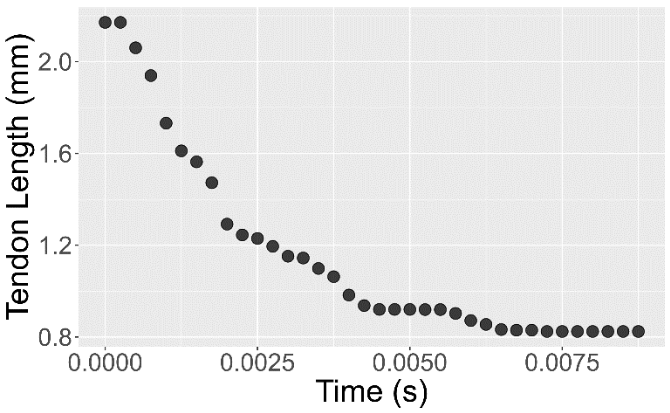

SEM Fig 4. Recoiling of the sternohyoideus tendon throughout the duration of a feeding strike. In this example the tendon shortens by 64% or 1.4 mm in 0.00475 s. Force as a function of length is known from tensile tests. This is then multiplied by the force of the tendon at equal length to the tendon at t =0 in order to obtain the power released from the tendon during the feeding event.

**Supplementary Methods**


26. S. W. Day, T. E. Higham, A. Y. Cheer, P. C. Wainwright, Spatial and temporal patterns of water flow generated by suction-feeding bluegill sunfish *Lepomis macrochirus* resolved by Particle Image Velocimetry. *J. Exp. Biol.* **208**, 2661–2671 (2005).
27. E. J. Stamhuis, Basics and principles of particle image velocimetry (PIV) for mapping biogenic and biologically relevant flows. *Aquat. Ecol.* **40**, 463–479 (2006).
28. Z. J. Taylor, R. Gurka, A. Liberzon, in *Handbook of Imaging in Biological Mechanics*, C. P. Neu, G. M. Genin, Eds. (CRC Press, 2014), pp. 173–184.
29. R. A. Holzman, D. C. Collar, S. W. Day, K. L. Bishop, P. C. Wainwright, Scaling of suction-induced flows in bluegill: morphological and kinematic predictors for the ontogeny of feeding performance. *J. Exp. Biol.* **211**, 2658–68 (2008).
30. K. L. Staab, L. A. Ferry, L. P. Hernandez, Comparative kinematics of cypriniform premaxillary protrusion. *Zoology*. **115**, 65–77 (2012).
31. J. K. Sveen, An introduction to MatPIV v. 1.6.1. *Mech. Appl. Math.* **2**, 1–27 (2004).
32. T. L. Hedrick, Software techniques for two- and three-dimensional kinematic measurements of biological and biomimetic systems. *Bioinspiration and Biomimetics*. **3**, 1–7 (2008).
33. C. E. Oufiero, R. A. Holzman, F. A. Young, P. C. Wainwright, New insights from serranid fishes on the role of trade-offs in suction-feeding diversification. *J. Exp. Biol.* **215**, 3845–3855 (2012).
34. T. E. Higham, S. W. Day, P. C. Wainwright, Multidimensional analysis of suction feeding performance in fishes: fluid speed, acceleration, strike accuracy and the ingested volume of water. *J. Exp. Biol.* **209**, 2713–2725 (2006).
35. S. Yaniv, D. Elad, R. A. Holzman, Suction feeding across fish life stages: flow dynamics

from larvae to adults and implications for prey capture. *J. Exp. Biol.* **217**, 3748–3757 (2014).

36. R. A. Holzman, S. Perkol-Finkel, G. Zilman, Mexican blind cavefish use mouth suction to detect obstacles. *J. Exp. Biol.* **217**, 1955–1962 (2014).

37. T. E. Higham, S. W. Day, P. C. Wainwright, The pressures of suction feeding: the relation between buccal pressure and induced fluid speed in centrarchid fishes. *J. Exp. Biol.* **209**, 3281–3287 (2006).

38. H. M. Dutta, Osteology, myology and feeding mechanism of *Astronotus ocellatus* (Pisces, Perciformes). *Zoomorphology*. **106**, 369–381 (1987).

39. G. van den Thillart, H. Smit, Carbohydrate metabolism of goldfish ( *Carassius auratus L* .). *J. Comp. Physiol. B*, 477–486 (1984).
